# Long-read cDNA sequencing reveals novel isoforms and spliceosome-mutant-enriched transcripts in AML and MDS

**DOI:** 10.64898/2026.05.20.726635

**Authors:** Christopher A. Miller, Sridhar Nonavinkere Srivatsan, Michael H. Kramer, Sai Mukund Ramakrishnan, Catrina C. Fronick, Robert S. Fulton, Casey D. Katerndahl, Nichole M. Helton, Timothy J. Ley, Matthew J. Walter

## Abstract

The alternative splicing landscape of the leukemia transcriptome remains poorly characterized, since short-read sequencing cannot resolve complete transcript structures. Using the Oxford Nanopore cDNA platform, we generated nearly 2 billion long reads (median 25.8 million per sample) from 71 human samples, including 48 acute myeloid leukemia or myelodysplastic syndrome samples, 25 of which had splicing-factor gene mutations (in *SRSF2, U2AF1*, or *SF3B1*). An additional 23 samples were from sorted hematopoietic cell populations from healthy individuals. Using this dataset, we created a transcript assembly containing 174,162 novel isoforms that are not described in the reference transcriptome. Deep-scale proteomic validation confirmed that many of these transcripts are translated into protein. We also identified isoforms enriched in spliceosome-mutant samples and found proteomic evidence of frequent nonsense-mediated decay regulation of novel transcripts. This dataset is a valuable community resource, enabling detection of new transcripts in short-read data sets. An interactive portal to explore splicing patterns in these data is available at https://leylab.org/isoforms/.

**Key Points:** Long-read sequencing enables the detection of many novel transcripts in AML, including many from splice-factor-mutant patient samples

This expanded transcriptome is a valuable community resource and can be used to improve analyses of short-read RNAseq data

## Introduction

Surveys of cancer cell transcriptomes have been undertaken in nearly every solid tumor and hematologic malignancy^1^. These studies have predominantly employed short-read cDNA sequencing on the Illumina platform, which provides deep coverage of individual exons and splice junctions, but is limited in its ability to link these elements together into complete transcript structures^2^. Long-read sequencing on the Pacific Biosciences or Oxford Nanopore (ONT) platforms can now generate reads that span cDNAs from end-to-end, but cost typically limits these experiments to lower read depths, ranging from 0.5 million reads to ∼10M reads per sample^3–7^. These efforts have revealed previously unknown splicing patterns and transcripts, but thus far, each increase in long-read sequencing depth continues to reveal new transcripts, suggesting that current datasets capture only a small fraction of the overall transcriptional complexity present in cancer^8^.

The issue of transcriptional representation is especially salient in cancers with mutations in spliceosome component genes, which are frequently observed in myeloid malignancies such as myelodysplastic syndromes (MDS) and acute myeloid leukemia (AML). In AML, the most common splicing-factor mutation is *SRSF2*, found in around 7% of cases, while *U2AF1* and *SF3B1* are each mutated in about 3%^9^. These mutations are far more common in MDS patients, with ∼50% harboring at least one mutation in a spliceosome gene^10–15^.

The splicing factor mutations assessed in this study impact either exon or splice-site recognition through distinct mechanisms, with effects ranging from dramatic to subtle. Well-characterized examples include “poison exon” creation, induced in *EZH2* (by *SRSF2* mutations)^18^ or *EIF4A* (by *U2AF1* S34F mutations)^19^; these create premature termination codons, induce nonsense-mediated decay and reduce transcript and/or protein levels. Cryptic splice site usage is another such example, where SF3B1 mutations lead to reduced *MAP3K7* levels^20^ and tumor progression. Previous studies have described thousands of splice alterations induced by these mutations (some using long-read sequencing at lower depths in other cancers)^5,7,21–23^. Determining which changes directly contribute to oncogenic phenotypes is an active area of investigation, but depends on first having high-quality full-length transcripts.

Short-read computational methods for detecting these changes (junction-by-junction) rely upon having a comprehensive and representative transcriptome to guide alignments and abundance estimation. Previous long-read sequencing in AML and MDS suggests that existing transcriptome assemblies are incomplete, and that thousands of novel transcripts may exist.

However, these studies used either low sequencing depths (∼0.5 million reads per sample)^22,24^ or moderate depths without splicing-factor mutations^4^. Given this low coverage and the known ability of spliceosome mutations to create novel transcripts, prior studies have underestimated the total transcriptional diversity of AML.

In this study, we paired deep long-read sequencing (median 25.8 million reads per sample) with a focus on splicing-factor mutations to search for transcriptional diversity absent from our current AML/MDS reference transcriptomes. We also included sorted populations from healthy donor bone marrow samples, including CD34-positive progenitor cells, relevant for clonal hematopoiesis and leukemogenesis, and CD3 or CD19-positive lymphocytes as references for immune cells, especially in a cancer context. Together, this approach reveals extensive and previously unrecognized isoform diversity in both myeloid malignancies and normal populations. This dataset will increase the power of the many existing short-read RNA sequencing studies. Coupled with an easy-to-use web interface, it will enable focused explorations of the role of transcriptome dysregulation in cancer.

## Methods

### Sequence generation and transcript assembly

Long reads were generated using the PCB111 kit on a PromethION machine, followed by trimming with pychopper, alignment to GRCh38 with minimap^26^, and statistics generation with Nanoplot^27^. Short reads were aligned with STAR version 2.7.0^28^ then quantified using Stringtie 1.3.3^29^ and/or kallisto v0.43.1^30^. Transcripts were assembled using ESPRESSO^31^ and filtered with SQUANTI3^33^.

### Key Analyses

Proteomic data was generated and processed as described previously^36^. FragPipe was used to re-search the data against novel protein sequences^37^. Enrichment statistics were calculated with custom R scripts (deposited for review and re-use). Differentially expressed isoforms were detected using rMATS-long v2.0.1^38^.

### Data Portal

The data exploration portal was based on IsoVis^39^, with substantial modifications made to enable abundance plotting, optimization for larger datasets, exposing additional transcript information, and accessory scripts for preparing data. The resulting code is available at https://github.com/chrisamiller/aml-transcriptome Detailed methodological descriptions are presented in the Supplemental Methods.

## Results

### Long-read cDNA sequencing from AML and MDS samples

We assembled a cohort of 71 human samples, from 48 patients with AML or MDS and 23 healthy donors (Table 1, Figure 1A, supplemental Table 1). 25 of these samples had hotspot mutations in spliceosome component genes: 16 with *SRSF2* P95[R/L/H] or G93GR, 5 with *U2AF1* S34[F/Y], 2 with *U2AF1* Q157P, and 2 with *SF3B1* K700E. As controls, we sequenced 23 samples from sorted hematopoietic cell populations from healthy individuals, including CD3+, CD19+, CD34+, promyelocytes (“Pros”), monocytes, and polymorphonuclear leukocytes (“PMNs”). All samples were sequenced on a PromethION device, producing a median of 25.8 million reads per sample (range: 16.6 - 69.3M), and 1.98 billion reads combined, for a total of 18.8 billion basepairs of coverage (Figure 1B, supplemental Table 1). Each sample was also sequenced with Illumina Total RNA short reads, producing a median of 135.1 million 2×151 bp reads per sample and 12.2 billion total basepairs.

**Table 1:** The patients in the cohort represent many subtypes of AML/MDS and are enriched for spliceosome mutations.

|  |  |
| --- | --- |
| **Age (mean +/- SD)** | 63.5 +/- 17.9 |
| **Sex (n, %)** |  |
| Male | 33 (68.7%) |
| Female | 15 (31.2%) |
| **Diagnosis (n, %)** |  |
| MDS | 3 (6.25%) |
| Primary AML | 42 (87.5%) |
| Secondary AML | 3 (6.25%) |
| **Key Mutations (n, %)** |  |
| *SRSF2* | 16 (33.3%) |
| *U2AF1* | 7 (14.5%) |
| *SF3B1* | 2 (4.2%) |
| *DNMT3A* | 7 (14.5%) |
| *FLT3* | 4 (8.3%) |
| *RUNX1* | 2 (4.2%) |
| *TP53* | 1 (2.08%) |
| **FAB Types** |  |
| M0 | 5 (10.4%) |
| M1 | 14 (29.2%) |
| M2 | 12 (25%) |
| M3 | 1 (2.1%) |
| M4 | 6 (12.5%) |
| M5 | 3 (6.2%) |
| unknown/NA | 7 (14.5%) |

**Figure 1:**
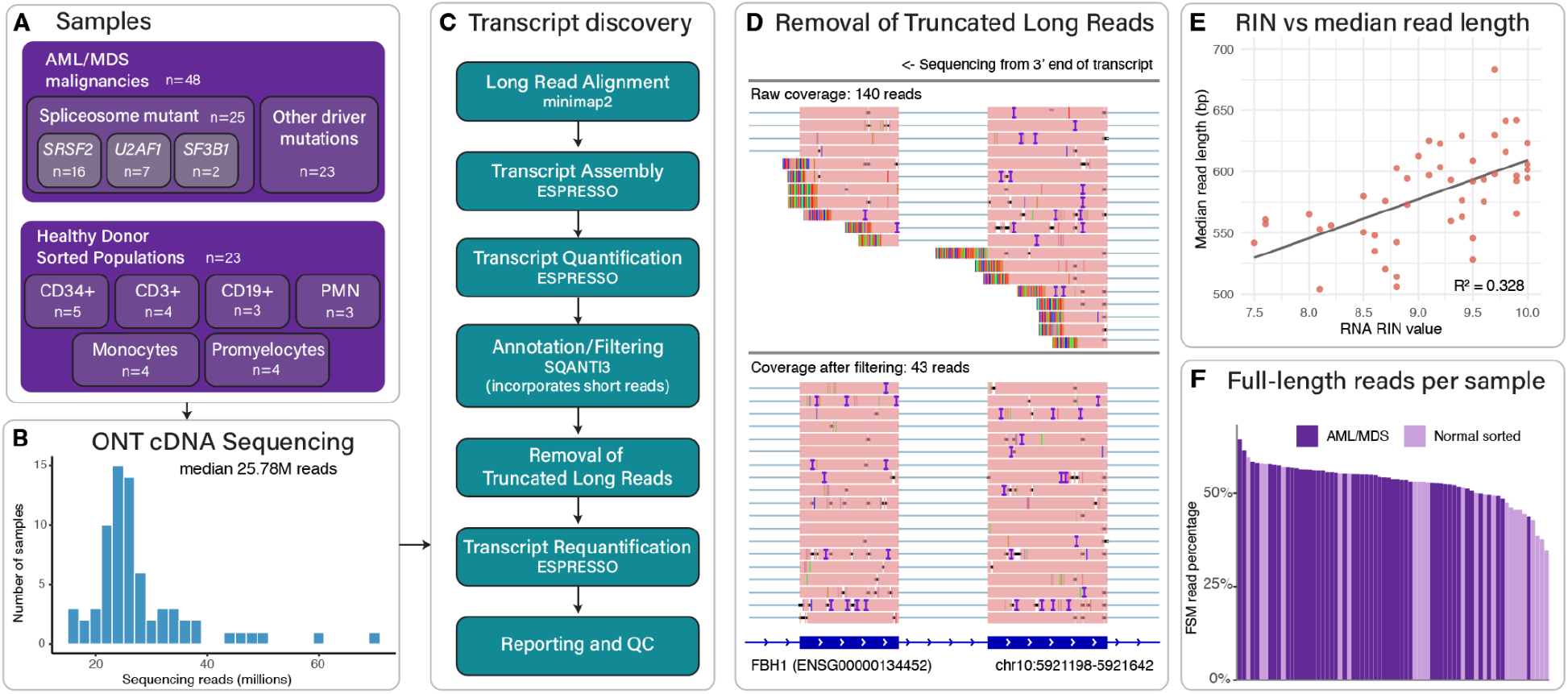
cDNA from AML/MDS and flow-sorted healthy normal samples was collected, then assembled and run through strict quality control and filtering. (A) Overview of the samples used in this cohort (B) Number of ONT reads generated per sample (C) Overview of the transcript assembly pipeline - see Methods for details (D) Representative region in the *FBH1* gene illustrating many reads with 5’ truncation artifacts that were removed by filtering. (E) Samples with lower RIN values tend to have shorter overall reads, as measured by median read length (Pearson’s correlation coefficient p=2.11×10^−5^) (F) Percentage of non-truncated, full-splice match reads in each sample.

### A scalable pipeline for artifact-aware long-read transcript assembly

We assembled the nearly 2 billion reads with ESPRESSO^31^, chosen in part because some other algorithms did not support straightforward parallelization that would accommodate this quantity of data^40,41^. Examination of the first-pass assembly revealed a substantial number of shortened isoforms, driven by striking 3’ bias and 5’ truncation (Figure 1D). These arose from truncated reads that have previously been attributed to homopolymer tracts^42^ (an association we could not replicate, perhaps due to platform improvements), reverse-transcriptase stalling^43^, or poor processivity of the pore enzyme or voltage spikes^44,45^. We observed a modest association between truncation hotspots and higher GC content (supplemental Figure 1), but this was not sufficient to explain the majority of the shortened reads. Notably, RNA Integrity Number (RIN) scores correlated with median ONT read length (Figure 1E), suggesting that optimizing for RNA quality is essential for reducing truncation artifacts.

Because ESPRESSO does not explicitly exclude truncated reads, we devised an iterative process to identify and exclude them (Figure 1C, Methods). Briefly, we assembled transcripts with ESPRESSO, then annotated and filtered the resulting transcripts with SQANTI3^33^. Key variables discriminating artifacts included read coverage, “bite” (signature of RNA secondary-structure artifacts), and TSS-ratio (comparing short to long-read coverage at transcript 5’ ends to flag likely truncations). This approach removed 368,343 putative transcripts.

The corresponding truncated ONT reads were then removed, eliminating 47.2% of all reads (median per sample 53.8%, range 34.8 - 64.5%, Figure 1F), an approach designed to prioritize accurate transcript assembly. More reads were discarded from fresh flow-sorted healthy normal samples than from cryopreserved tumor samples (mean 54.5% vs 49.5%, Wilcoxon p=0.002). After requantification, we required novel isoforms to be called in at least 2 samples and supported by 5 total long reads. Since long transcripts were underrepresented, we also retained all Ensembl transcripts, reducing false negatives while focusing our discovery efforts on novel transcripts, especially those caused by aberrant splicing^32^.

### Novel transcripts expand the reference transcriptome

Despite this conservative approach, the resulting transcriptome assembly contained 174,162 novel transcripts, along with 206,601 known Ensembl transcripts (Figure 2A). Of these novel transcripts, 51.2% were identified in at least 10 samples, with 1.5% detected in all samples (Figure 2B). Novel transcripts had lengths comparable to known Ensembl transcripts (Figure 2C), though a smaller fraction exceeded 3,000 nucleotides in length, reflecting the 3’ bias in the data (known: 17.9%, novel: 11.0%, Chi-squared *p* < 2×10^−16^). Novel transcripts exhibited lower overall expression (Figure 2D) and collectively represent 3.7% of the total mRNA quantified (Figure 2E), consistent with the expectation that highly expressed isoforms were historically easier to detect. Nonetheless, 10,186 transcripts contribute over 10% of their corresponding gene’s total expression. Together with extensive manual review, these characteristics gave us high confidence that the transcripts reported here are predominantly genuine, novel isoforms.

**Figure 2:**
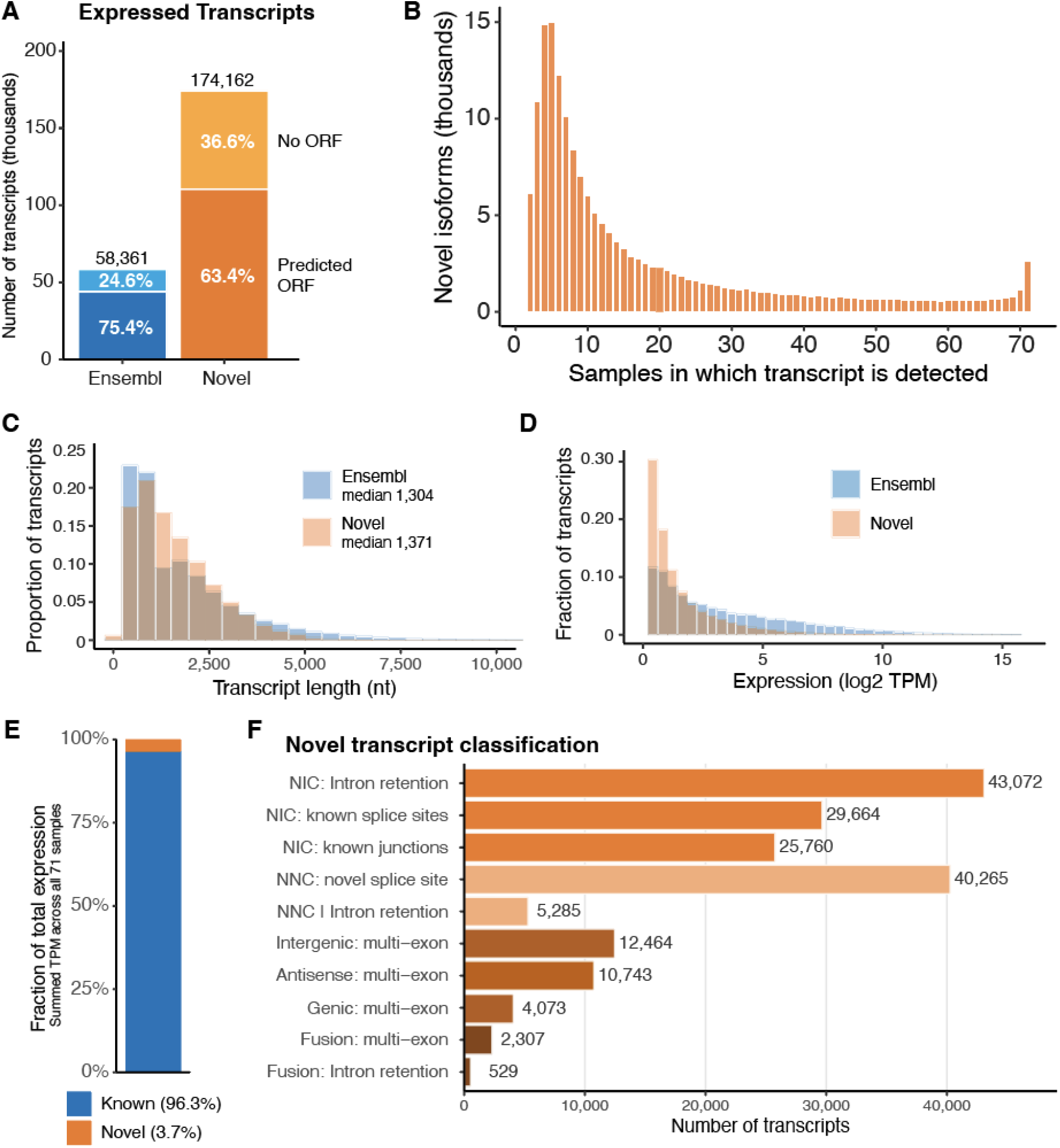
Transcript assembly reveals high quality novel isoforms in this dataset. (A) Transcripts with measurable expression in this cohort (only 58,361 of 206,601 Ensembl transcripts were detected in these tissues). Significantly fewer novel transcripts are predicted to have open reading frames (ORFs). (Fisher’s exact test, *p*<2×10^−16^). (B) The number of samples expressing each novel transcript (C) Distribution of transcript lengths, comparing known Ensembl to newly identified transcripts (D) Distribution of total expression levels (TPM) for known and novel transcripts, summed across all samples (E) Fraction of total expression that comes from known vs new transcripts (F) Classification of novel transcripts from SQANTI3. (NIC = Novel In Catalog, NNC = Novel Not in Catalog)

Intron retention was the most common class of novel transcript (Figure 2F). A plurality of the rest were made up of novel splice sites and combinations of known splice sites and junctions, categories that include traditional short-read classifications, including exon skipping or alternate 3’ or 5’ splice sites. 23,207 of these transcripts (13.3%) came from 7,048 novel regions mapping entirely outside known annotations. Of these, 992 intersected with known long non-coding RNAs^34^, and another small fraction represented partial overlaps to sequences not represented in Ensembl (predicted genes, other small RNA species, or antisense transcripts), but the majority represented expression from poorly annotated sequences.

### Proteomic evidence for translation of novel transcripts

24 of the AML samples in this study had previously-generated deep-scale proteomic data, enabling a search for evidence of novel isoform translation^36^. Several factors make such identification challenging: 1) The relatively low sensitivity of mass spectrometry (this “deep” proteome detected a mean of only 8,976 proteins per sample), 2) Low expression of most novel isoforms, implying that their protein abundance may also be low, 3) variable peptide ionization, and 4) the requirement that novel events generate a detectable tryptic peptide. Despite these limitations, we identified 307 distinct high-confidence peptides from 273 different transcripts and 261 genes (Figure 3A, supplemental Table 2). These fall into several categories, with 160 (52.1%) representing a new mRNA sequence derived from known genes -- generated by intron retention, alternative 3’ or 5’ splice sites, frameshifts, or use of new exons. Another 126 (41.0%) were fusion peptides from splicing alterations (e.g. exon skipping). Eight (2.6%) originated from novel protein coding sequences, and 13 (4.2%) were from gene fusion events, primarily read-through transcripts from neighboring genes. Figures 3B-D provide representative examples. Although many of these may generate neoantigens that could theoretically be recognized by the immune system, the low overall expression of these transcripts and peptides may limit their attractiveness as therapeutic targets.

**Figure 3:**
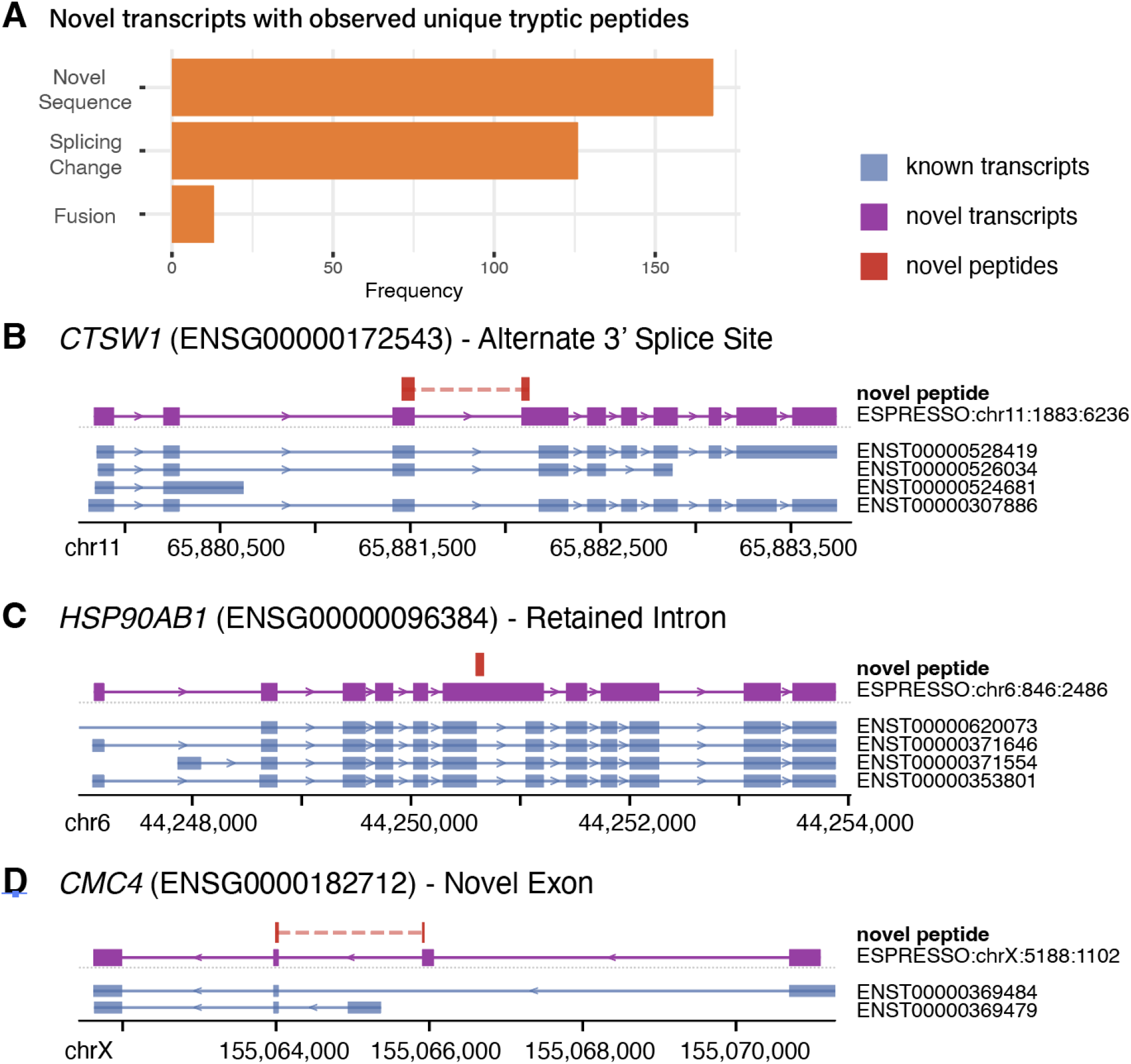
Novel transcript translation was validated with deep-scale proteomics. (A) Classification of novel peptides detected in any of 24 AML samples evaluated with TMT proteomics (B) Schematic of the *CTSW* gene, showing known transcripts, novel transcripts, and the location of the translated sequence that gives rise to a novel peptide, in this case from sequence spanning the junction between a known exon and new sequence created from alternate 3’ splice site usage (C) Schematic of the *HSP90AB1* gene, showing a novel peptide translated from retained intron sequence (D) Schematic of *CMC4*, which has a novel peptide created from sequence that spans a splice junction from a known exon into a novel exon.

Standard protein database searches often use “reviewed” UniProt references, which provide a less comprehensive set of protein sequences than those described in either Ensembl or our expanded assembly^46^. Searching against peptides predicted from our extended transcriptome identified thousands of additional peptide matches, falling largely into two categories: fully tryptic peptides in known Ensembl transcripts, but absent from UniProt, and semi-tryptic partial matches potentially arising from alternative splicing that creates premature termination codons or alternative start sites. Since we could not rule out protein degradation or non-specific proteolytic cleavage as a cause of these truncated peptide observations, we did not treat them as strong evidence for novel isoform translation. Collectively, these analyses suggest that more comprehensive reference databases may enhance proteomics studies and they also provide evidence that many novel transcripts are valid and being translated into proteins.

### Characterization of nonsense-mediated decay in novel transcripts

The majority (63.4%) of novel transcripts are predicted to have protein-coding ORFs, but show significantly higher levels of predicted nonsense-mediated decay (NMD) than known transcripts (*p* < 2.2×10^−16^, Figure 4A), a finding consistent with previous reports^47^. While tumors and healthy cells have similar overall levels of predicted NMD transcript expression, a larger proportion of this expression comes from novel isoforms in tumor samples (Figure 4B-C, *p*=4.68×10^−7^), suggesting that the contribution of NMD may have previously been underestimated in tumor cells. Overall, 417 genes had significantly higher levels of predicted NMD transcript accumulation in the AML/MDS samples than in healthy cells, and 337 genes had elevated predicted NMD specifically in splicing factor-mutant tumor samples (supplemental Tables 3, 4). Both lists had overrepresentation of genes in RNA metabolism and mRNA splicing (largely driven by novel transcripts) supporting that NMD may be a common self-regulatory mechanism for these pathways^48–51^.

**Figure 4:**
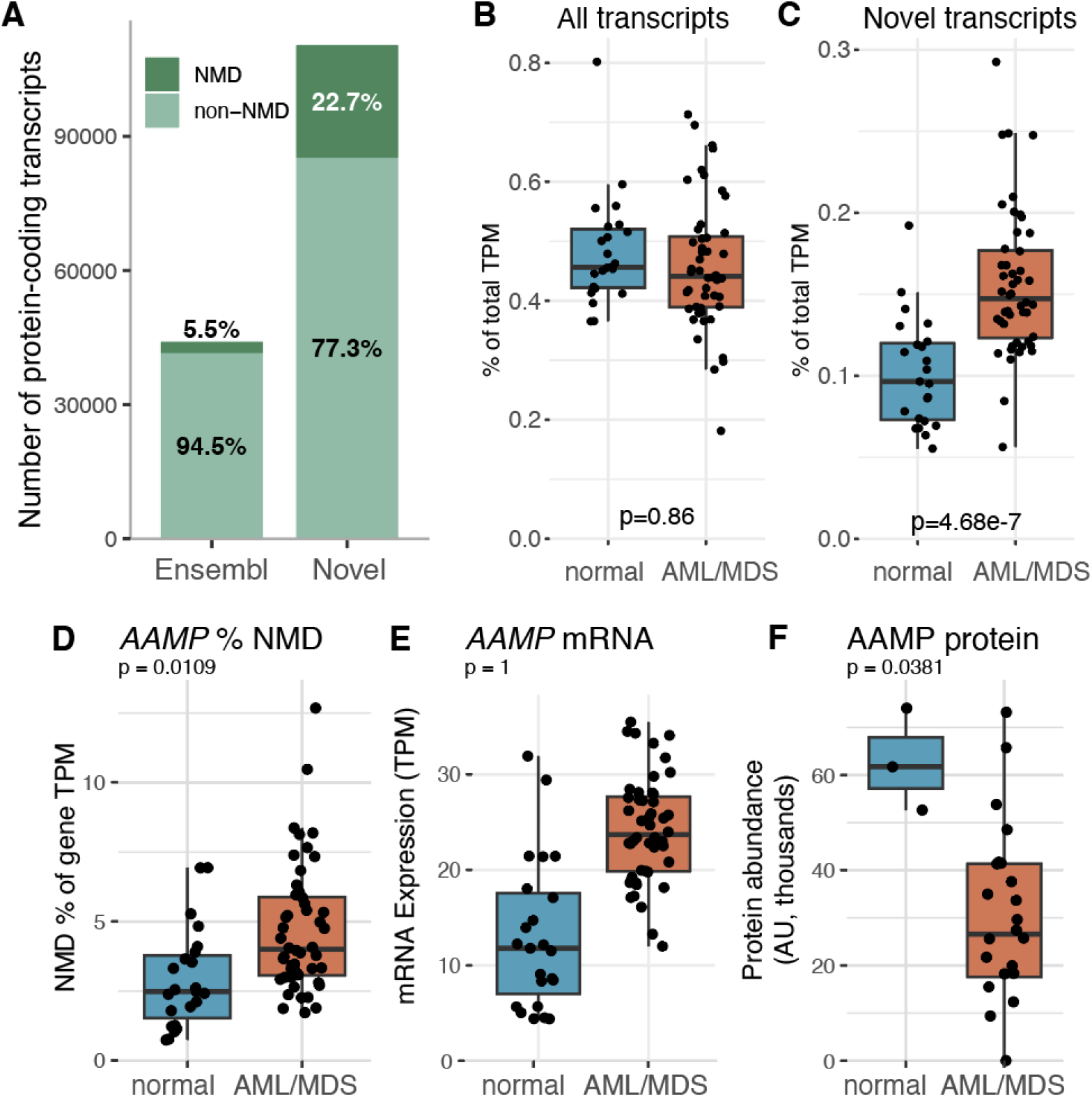
Nonsense-mediated decay is increased in novel transcripts. (A) Number and percentage of predicted protein-coding transcripts predicted to undergo nonsense-mediated decay (NMD) (Fisher’s exact test: *p*<2.2×10^−16^). (B) Percentage of all mRNA expression that comes from NMD transcripts in normal healthy cells (blue) and AML/MDS tumor cells (red) (Wilcoxon rank sum *p*=0.86). (C) Percentage of all mRNA expression that comes from novel NMD transcripts in normal healthy cells (blue) and AML/MDS tumor cells (red) (Wilcoxon rank sum, *p*=4.68×10^−7^). (D) Increased percentage of overall *AAMP* transcripts coming from NMD transcripts in tumors as compared to normal cell populations (Wilcoxon rank sum, *p* = 0.0109). (E) mRNA expression of *AAMP* in tumors compared to normal healthy cells (F) Protein abundance of AAMP in tumors compared to normal cell populations (Wilcoxon rank sum, *p*=0.0381) All *p* values BH-adjusted.

NMD regulation can occur via rapid mRNA degradation, or with more stable mRNAs that have altered translation or that produce proteins with short half lives^52^. By integrating the proteomics data above with our expanded transcriptome, we searched for genes subject to NMD-mediated regulation that would be invisible without both data layers. We specifically looked for genes where: 1) gene-level mRNA expression was not significantly reduced in tumors, 2) predicted NMD transcripts made up an increased proportion of the gene’s expression, and 3) protein abundance was significantly lower in tumor samples (all using Wilcoxon rank sum test,BH-corrected *p*<0.1). We identified 70 such genes, including *AAMP* (Figure 4D-F), where the NMD isoform shift occurs primarily within novel transcripts. This NMD shift is not unique: across tumor samples, a mean of 63% of all NMD transcripts were novel (range 37-80%). The large number of newly detectable predicted NMD transcripts in this assembly will facilitate more detailed and accurate characterization of these effects in future studies, especially when coupled with proteomics.

### Healthy donor cells harbor population-specific novel transcripts

Using data from sorted populations of bone marrow cells from healthy donors, we sought to identify novel isoforms that were restricted to or enriched in specific hematopoietic lineages and/or differentiation states (Figure 5A-B, supplemental Figure 2, supplemental Table 5). Each sorted cell type contained high numbers of such events, ranging from 850 in CD19-positive B cells to 9,580 in monocytes. AML/MDS cells harbored more specific/enriched novel transcripts (21,101), perhaps due to their altered biology and the greater statistical power generated by a larger number of samples. Many of these population-enriched novel transcripts occurred in well-known cell lineage regulated genes, like *TCF7* in CD3-positive T cells (Figure 5B), or *FAM133A* in CD34-positive myeloid progenitor cells (Figure 5C), revealing new splicing patterns in these genes. Overrepresentation analysis confirmed that genes bearing these novel transcripts are significantly associated with known lineage-relevant pathways, like “T Cell Activation” and “Lymphocyte Differentiation” for CD3+ cells (supplemental Table 6).

**Figure 5:**
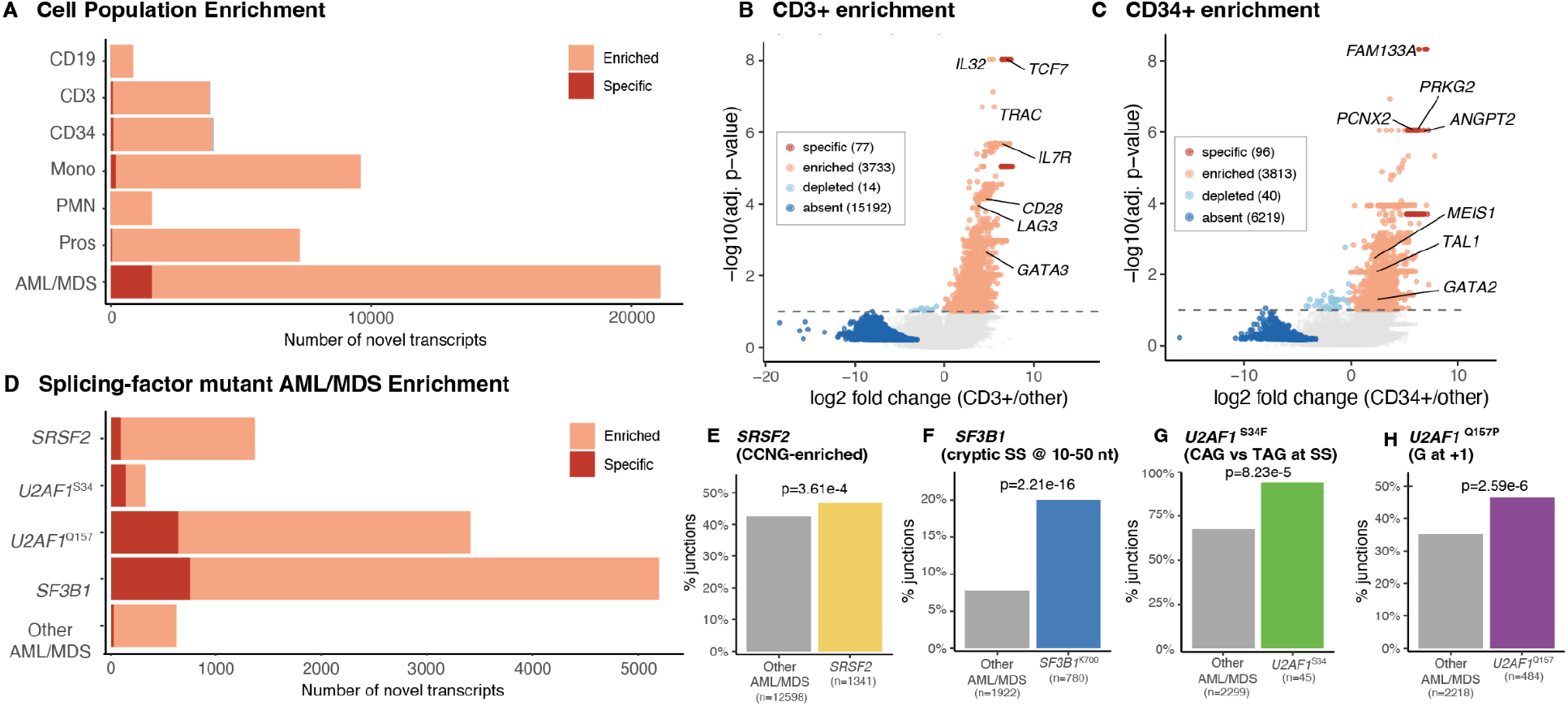
Subsets of samples are enriched for specific novel transcripts and signatures. (A) Quantification of novel transcripts that are enriched in (pink) or specific to (red) sorted populations or tumor cells, as compared to all other samples combined (B) novel transcript enrichment in CD3+ sorted T cells (C) novel transcript enrichment in CD34+ sorted stem/progenitor cells (D) Counts of novel transcripts that are enriched (pink) or specific to (red) splicing-factor mutant AML/MDS samples, as compared to all other AML/MDS and CD34+ stem/progenitor cells. (E) Enrichment of CCNG motifs (vs GGNG) in novel transcripts from *SRSF2*-mutant samples (Wilcoxon rank sum *p*=3.61×10^−4^) (F) Fraction of alternative 3’ splice site junctions that occur in the expected interval (10-50bp upstream of a known acceptor site) in SF3B1 samples (Fisher’s exact test *p*=2.21×10^−16^) (G) Fraction of alternative 3’ splice site junctions that match the CAG motif vs TAG motif in samples with *U2AF1*^S34F^ mutations (Fisher’s exact test *p*=8.23×10^−5^). (H) Fraction of alternative 3’ splice site junctions that have G at position -1 vs C in samples with *U2AF1*^Q157P^ mutations (Fisher’s exact test *p*=2.59×10^−6^)

**Figure 6:**
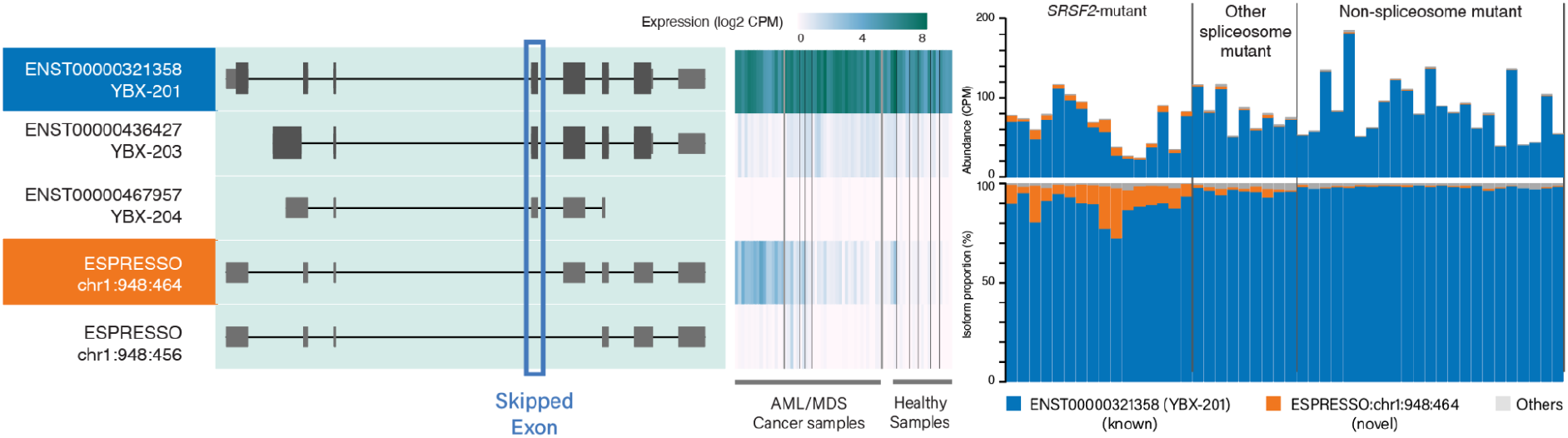
*YBX1* harbors a novel transcript with high abundance in splicing-factor-mutated samples. Exon skipping results in a novel transcript of *YBX1* (left), that has increased abundance in *SRSF2*-mutant and *U2AF1*-mutant AML/MDS samples (center and right).

### Spliceosome mutations are associated with characteristic novel transcripts

The splicing-factor gene mutations in our cohort are known to cause specific, well-defined alterations in splice site recognition and usage. We therefore hypothesized that novel transcripts enriched in splicing-factor-mutant tumors would bear the sequence signatures characteristic of the respective mutant splicing factor. We performed the same enrichment analysis described above on the AML/MDS cohort, stratifying samples by the presence of mutations in *SRSF2, SF3B1*, and *U2AF1* (separating S34 vs. Q157 mutations, since they are known to have distinct functional effects^53^, Figure 5D, supplemental Figure 3, supplemental Table 7). We then compared these samples to non-splicing factor mutated AML/MDS samples to determine the proportion of the novel splicing events that contained the canonical splice site sequence motif associated with each mutation.

Novel transcripts enriched in *SRSF2-*mutant samples displayed the expected shift towards CCNG over GGNG motifs in exon splicing enhancers (*p*<2.1×10^−5^)^18,54^. Similarly, novel transcripts enriched in *SF3B1*-mutated samples showed preferential use of cryptic 3’ splice sites located 10-50 nucleotides upstream of a canonical acceptor site (*p*=2.7×10^−19^)^55^. In *U2AF1* S34F-mutant samples, cytosine at the -3 position, preceding the splice-acceptor AG dinucleotide, was significantly enriched relative to the wild-type thymine preference in novel transcripts (*p*=0.001). *U2AF1* Q157P mutant samples showed evidence of increased guanine residues at the +1 position relative to the 3’ splice site (*p*=2.1×10^−4^, Figure 3B)^56^. Overrepresentation analysis of novel transcript-harboring genes with these four splicing-factor gene mutations showed significant enrichment in pathways like “RNA splicing” and “mRNA processing” (supplemental Table 8), which support the fact that these factors are known to function in self-regulatory feedback loops^57,58^. *DNMT3A* inactivation in mouse hematopoietic cells has been recently postulated to cause altered mRNA splicing, but data from the 7 *DNMT3A*-mutated samples in this study did not show any evidence that would support the proposed mechanism^59^. Collectively, these results provide evidence that mutationally-driven splicing mechanisms contribute to novel transcript generation in AML/MDS.

### Detecting YBX1 loss-of-function splice variants in *SRSF2*-mutant samples

By applying rMATS-long to these data, we identified 1,524 transcripts that were differentially expressed between AML/MDS and healthy donor derived hematopoietic cells, and between 755 and 2,642 that were differentially expressed between specific population or tumor subsets (supplemental Table 9). Overall, 24% of differential isoforms were from novel transcripts. Among the most notable findings was a novel isoform of *YBX1*, a gene with proposed roles in AML pathogenesis^60–63^. This transcript (ESPRESSO:chr1:948:464) is nearly absent in most AML/MDS and healthy donor samples but represents up to 25% of YBX1 transcripts in *SRSF2*-mutant samples. It also has less pronounced overexpression in *U2AF1*-mutant, but not *SF3B1*-mutant, AML/MDS samples. This exon skipping event deletes much of the functional cold-shock domain, with predicted deleterious effects. Further work will be needed to understand the functional consequences of this isoform switch and whether it contributes to AML/MDS pathogenesis.

### A more complete transcriptome assembly enables new insights from short-read data

We next evaluated whether the expanded transcriptome could improve transcript quantification from short-read RNA-seq data. By realigning and requantifying the short-read RNAseq data from 65 of this study’s samples, we detected 95.2% (165,724/174,162) of the novel transcripts,none of which could be reported from the original short-read data processing. Importantly, these short-read sequences were not used to create this assembly directly - their sole prior use was as a coverage filter. More nuanced splice-junction analyses would likely recover some of these events, but using a more comprehensive transcriptome means that they are immediately included in the primary results.

We then extended this approach to 130 additional short-read RNA-seq samples, recovering 96.2% of novel transcripts (167,679/174,162, Figure 7A). The alternative *YBX1* transcript described above (ESPRESSO:chr1:948:464) was detected at high levels (>5 CPM) in 6 of those samples. When cross-referenced with DNA sequencing data, 5 of the 6 AMLs harbored *SRSF2* P95 mutations, and the other contained a *U2AF1* Q157P mutation, further validating the mutational association described above. Notably, *YBX1* had no differential expression at the gene level, so it would not have been flagged as a gene of interest by standard analyses.

**Figure 7:**
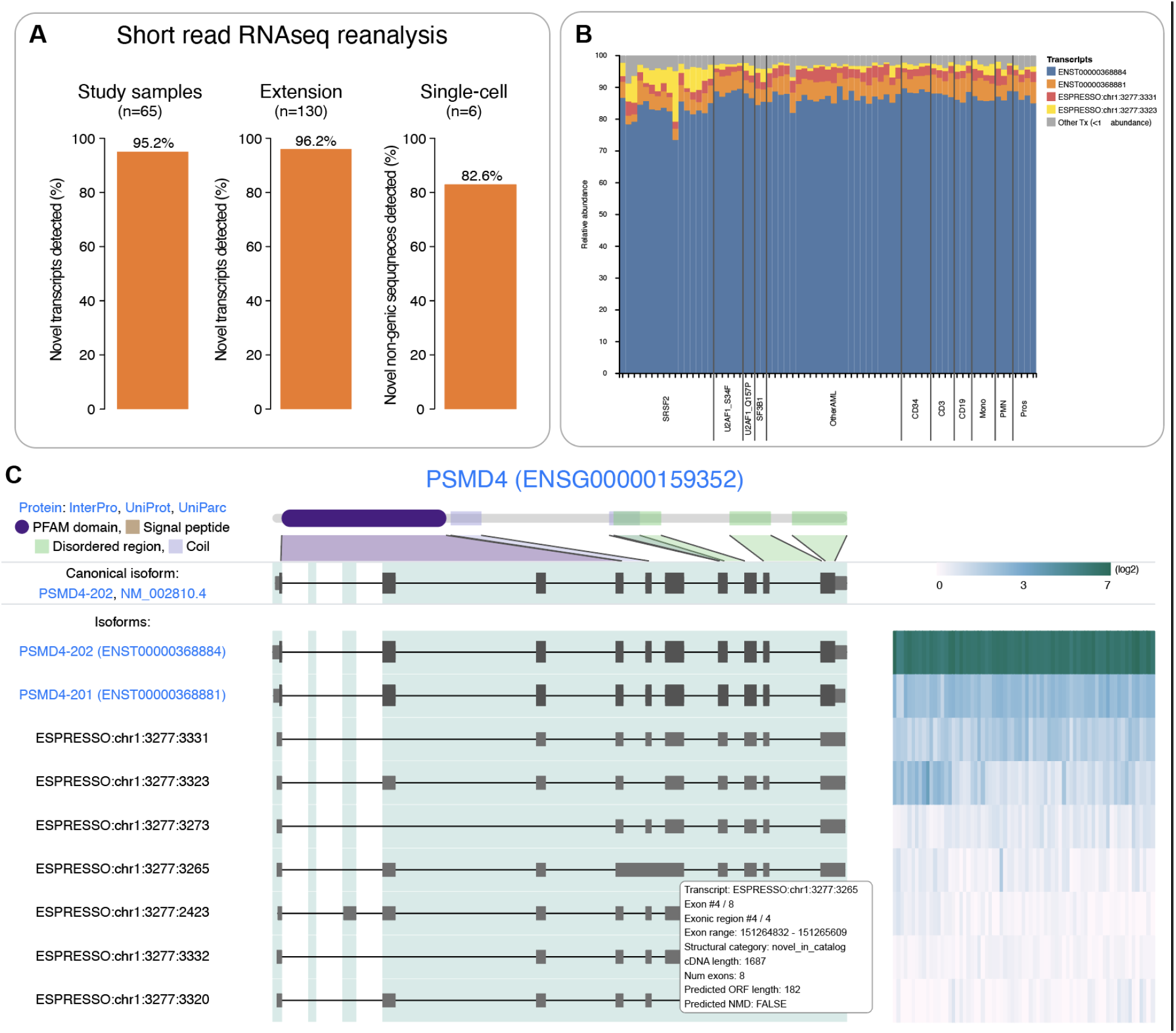
This resource allows for deep exploration of the expanded transcriptome. (A) Processing short reads with the expanded transcriptome enables primary detection of most novel isoforms, both in short-read data from this cohort (left) and in an extension short-read cohort (center). Novel non-genic sequences are similarly newly accessible in single-cell RNAseq data (right). (B) Screenshot of transcript abundance generated from the web portal (C) Screenshot of the web portal displaying transcript structures (left), abundance heatmaps (right), and detailed transcript information (popup). Additional visualization layers and filters are available through the portal’s menus.

Finally, we aligned single-cell RNAseq (5’ data kit from 10x Genomics) from 6 AML samples (not from this cohort) to this expanded reference. Since scRNA-seq pipelines aggregate to the gene level to improve sensitivity, we restricted our search to novel sequences (not overlapping known genes). In these samples, we detected transcripts from 5,829 of 7,049 (82.6%) such “genes”, and 18,362 out of 31,370 cells (58.5%) contained reads from one or more. These results underscore the fact that many transcripts are inaccessible using the standard Ensembl transcript reference, but are recoverable using this expanded reference transcriptome, even when using short-read data.

### A resource for improving transcriptomics in blood cancers

Along with this expanded transcriptome, we developed a “Transcriptome Explorer” web portal that provides detailed information on transcripts, splicing patterns, and gene and transcript abundance (Figure 7B/C) from this study. The portal was built on top of the IsoVis software, heavily modified and expanded to add functionality specific to this data set^39^. It is freely available for use at https://leylab.org/isoforms/. The source code can be found at https://github.com/chrisamiller/aml-transcriptome. The portal also hosts key datasets and a GTF ready for use withRNAseq pipelines.

## Discussion

In this study, we sequenced long-read cDNAs at high depth from a broad set of bone marrow samples from healthy donors or patients with myeloid malignancies, creating a transcript assembly more comprehensive than the Ensembl reference transcriptome. The resulting catalog -- 174,162 novel transcripts alongside 206,601 known Ensembl isoforms -- represents 1.5-fold more novel transcripts than reported in a previous, lower sequencing depth study of AML (direct comparison to the work of Shi, et al., was not possible because their assembly is not publicly available)^24^. This expansion is primarily attributable to two factors: The sequencing depth (nearly 2 billion reads, median 25.8 million per sample), and the inclusion of 25 splicing-factor mutant samples, which are a particularly rich source of transcriptome diversity and have not previously been deeply characterized with long reads.

Proteomic data contributed greatly to this study and pairing it with deep transcriptional profiling suggests a paradigm for future studies. Detection of 307 high-confidence novel peptides demonstrates that some novel isoforms are translated, and that larger transcript assemblies can aid in recovering biologically-relevant proteins that are currently ignored. We also identify genes like *AAMP* where protein levels are reduced despite no reduction in overall mRNA abundance. Isoform shifts towards NMD-sensitive transcripts suggest that NMD regulation is occurring but is invisible to gene-level analysis. These findings emphasize that comprehensive transcript catalogs are necessary to fully characterize NMD regulation in cancer cells.

The inclusion of flow-sorted bone marrow subpopulations allowed us to deeply characterize cell types that comprise only a few percent of a typical bone marrow sample. These included CD3+ T cells, CD19+ B cells, and CD34+ stem/progenitor cells, which are often the cell of origin for MDS and AML. Each population contains hundreds to thousands of population-enriched novel transcripts, often in lineage-relevant genes, with potential utility for immune or functional studies in many tumor types.

Unlike prior work that characterized splicing-factor gene mutant AML/MDS at the level of individual splice junctions, this full-length assembly places aberrant junctions into the context of the entire transcript structure. Although we can detect the mutation-specific signatures of splicing dysregulation, fewer than 0.1% of novel transcripts in splicing-factor mutant samples are specific for a given genotype. Instead, these data support a model in which spliceosome mutations shift the relative ratios of canonical splice-sites, enriching for low-abundance isoforms, rather than creating entirely new ones^55,64^. This more comprehensive catalog of altered splicing in these samples will aid in detailed studies of specific genes and pathways that may be relevant for oncogenesis.

An illustration of this resource’s value is the identification of a novel *YBX1* exon-skipping isoform, highly enriched in *SRSF2*-mutant AML and MDS. Though nearly absent in other AMLs, and in healthy donor cells, this isoform represents up to 25% of *YBX1* transcripts in *SRSF2*-mutant samples. Validation using 130 short-read AML samples confirmed the association and identified one cryptic *SRSF2* mutation that was previously missed. *YBX1* is one of several strong candidates for functional follow-up studies and represents a broader class of spliceosome-driven isoform switches that do not alter overall gene expression, but can be readily detected at the transcript level by using a more comprehensive reference assembly.

This expanded transcriptome resource is immediately applicable to existing RNA-seq datasets. When reanalyzing short-read RNAseq samples we were able to detect most novel isoforms, and in a reanalysis of 6 single-cell RNAseq samples from AML patients, more than 50% of novel genes were recovered. Applying these same techniques to other large short-read datasets, like Beat AML or TCGA LAML, represents an immediate opportunity to discover previously overlooked isoform shifts that may be relevant for disease pathogenesis^65,66^.

Limitations of this study include that this assembly was based on v95 of Ensembl, rather than the latest release (v115), to maintain compatibility with other AML datasets. A post-hoc analysis found that 12.3% of our novel transcripts are now present in Ensembl 115, which reflects the increasing breadth of databases from studies like this one, and serves as further validation of the quality of this assembly. 3’ bias is perhaps the largest limitation of this study, because ONT sequencing with poly-A+ priming limits the representation of longer transcripts. Neither alternative platforms (Pacific Biosciences has similar 3’ issues) nor direct RNA sequencing (low yields) would solve this problem at this time. We observed that RNA integrity was also relevant for sequence quality, as is flow-sorting, which may compromise RNA quality even more than cryopreservation. These are all findings with practical implications for study design. Cohort size was also somewhat limiting, with small numbers of *SF3B1* mutant (n=2) and *U2AF1* Q157P mutant (n=2) samples and a restricted set of sorted cell populations from healthy donor bone marrows. Future studies could fill these gaps by expanding the repertoire of cell-types, including other spliceosome mutations (e.g. *ZRSR2*) and pursuing deeper proteomic coverage, which could potentially identify a larger number of peptides derived from novel isoforms.

In conclusion, this study represents the deepest long-read transcriptome of AML/MDS samples to date and reveals more extensive isoform diversity than previously described. The expanded transcript assembly is immediately applicable to existing and future short-read, single cell, and proteomic datasets, and the “Transcriptome Explorer” web portal makes these data accessible to the broader research community. Studies built upon this reference will be well-positioned to connect specific isoform shifts to clinical phenotypes, therapeutic approaches, and the mechanisms of spliceosome-driven oncogenesis.

## Supporting information

Supplemental Methods, Results, and Figures

## Acknowledgements

Supported by the National Cancer Institute (NCI)/NIH R50CA211782 (to C.A.M.), NCI grants P01CA101937, R35CA197561 (to TJL), SPORE in Leukemia P50CA171963 (Daniel Link, PI),an Edward P. Evans Foundation grant and Blood Cancer United (to M.J.W.), and a Barnes Jewish Hospital Foundation Award (to T.J.L.). Thanks to Katherine Melville, Maddy Hartley, Brandon Blakely, and Jerid Robinson from Oxford Nanopore Technologies for useful discussions and troubleshooting.

## Authorship Contributions

Conceptualization, C.A.M, S.N.S., T.J.L, M.J.W.; Methodology: C.A.M, S.N.S.; Software: C.A.M.;Formal analysis: C.A.M. S.N.S., S.M.R.; Investigation: C.A.M., S.N.S., M.H.K., C.C.F., R.S.F.,C.D.K., N.M.H.; Writing - original draft: C.A.M, M.J.W.; Writing - review and editing: C.A.M, T.J.L., M.J.W.; Visualization: C.A.M.; Supervision: C.A.M, T.J.L., M.J.W.; Funding Acquisition: C.A.M., M.J.W., T.J.L.

## Disclosure of Conflicts of Interest

No conflicts to report.

—-------------------------------

## SUPPLEMENTAL INFORMATION

**Supplemental Methods and Supplemental Figures 1-3:** see attached

**Supplemental Table 1:** Detailed characteristics of the patient cohort and ONT sequencing

**Supplemental Table 2:** High-confidence peptides detected from novel transcripts

**Supplemental Table 3:** Genes with higher levels of NMD transcript accumulation in tumor samples

**Supplemental Table 4:** Genes with higher levels of NMD transcript accumulation in splicing-factor mutant tumor samples

**Supplemental Table 5:** Novel transcripts specific to or enriched in a given population

**Supplemental Table 6:** Overrepresentation analysis of genes harboring enriched novel transcripts in specific populations

**Supplemental Table 7:** Novel transcripts specific to or enriched in categories of splicing-factor-mutant tumors

**Supplemental Table 8:** Overrepresentation analysis of genes harboring enriched novel transcripts in splicing-factor mutant tumor samples

**Supplemental Table 9:** Differentially expressed isoforms between specific population or tumor subsets from rMATS-long

