## Supplemental Methods, Results, and Figures for "Long-read cDNA sequencing reveals novel isoforms and spliceosome-mutant-enriched transcripts in AML and MDS"

### Supplementary Methods

#### Sequence generation

For each sample, the quality of the total RNA was determined by the Agilent Bioanalyzer (RIN ranged from 6.7-10), the quantity was determined by the Qubit Flex (HS RNA kit), and the purity was assessed on the nanodrop (expected values 260/280 ~2 and 260/230 ~2-2.2). 200ng of total RNA was input into the PCR-cDNA Barcoding Kit (Oxford Nanopore, SQK-PCB111.24). The reverse-transcribed (RT) sample was split into two second strand PCR reactions (2x50uL reactions with 5uL of RT product in each reaction per the manufacturer's protocol). 14 cycles of PCR with 8-minute extensions were performed. Unique barcoded adaptors were ligated per sample. The barcoded libraries were pooled in equal molar ratios and loaded across multiple flow cells to mitigate flow cell to flow cell pore count variability. ~50fmol of pooled library was loaded per PromethION flow cell (R9.4.1, Oxford Nanopore, FLO-PRO002) targeting 30-40M reads per sample. Sequence data was base called on-instrument using the Super-accurate basecalling model (Guppy 6.4.6). Data was collected for ~90 hours generating >140M reads per flow cell. Short read RNA seq was prepared using the TruSeq Stranded Total RNA Gold kit (Illumina RS-122-2303), followed by sequencing on an Illumina machine.

#### Data Processing

Long reads were trimmed with pychopper (https://github.com/epi2me-labs/pychopper), then aligned to human genome GRCh38 with the chromosome 21 U2AF1 patch (https://ftp.ncbi.nlm.nih.gov/genomes/all/GCA/000/001/405/GCA_000001405.15_GRCh38/seqs_for_alignment_pipelines.ucsc_ids/GCA_000001405.15_GRCh38_GRC_exclusions.bed) using minimap2 v2.17 with parameters `-ax splice -uf`. Quality control statistics were generated using NanoPlot version 1.41.0. Short reads were aligned with STAR version 2.7.0 then quantified using Stringtie 1.3.3 and/or kallisto v0.43.1.

#### Transcript Assembly and filtering

We first ran the standard ESPRESSO workflow on the aligned reads, using version 1.3.2, parallelized per-chromosome in order to facilitate lower memory usage and increased speed. Ensembl version 95 transcripts were provided as a baseline reference. Annotation of the assembled transcripts was performed with SQANTI3 version 5.2.1, providing the short-read RNAseq to provide evidence for specific splice junctions and coverage patterns that support the assembled long-read transcript models. SQANTI3 was then used to filter the data, using the built-in machine learning approach with default parameters. Novel transcripts that passed these filters were merged with the original Ensembl transcripts. ESPRESSO quantification was then re-run so that reads linked to removed transcripts had a chance to be reassigned. The read assignments from this step were processed and all reads not categorized as full-splice-match are removed as probable truncation artifacts. Quantification was run a third time on these full-splice-match reads to produce a final long-read expression matrix. Coverage filters were then applied, to remove any transcript not present in at least two samples and not supported by at least 5 reads. Comparisons of this assembly to other ensembl versions or to LNCipedia were carried out using gffcompare 0.12.6.

**Proteomics**

Proteomic data was generated and processed as described previously, and abundance values from that publication were used in all plots, unless otherwise specified. For detection of novel peptides, open reading frame predictions from SQANTI3 were used to identify peptides unique to novel transcripts. FragPipe version 22.0 was used to re-search the TMT data against this set of novel protein sequences using the default TMT-11 workflow parameters (using trypsin digest and allowing 2 missed cleavages). The resulting protein abundance files were compared to all peptides from Ensembl v95 to remove known sequences. Each peptide was then classified according to its mechanism of creation (splicing, fusion, frameshift, etc).

**Transcript Enrichment and NMD**

For population-based enrichment, each sorted normal population was grouped individually with tumors combined as a separate group. Transcripts were tested for enrichment/depletion in a particular population using a two-sided Wilcoxon rank-sum test. The "specific" label was applied when a transcript was only found in one group, regardless of significance. The "absent" label similarly required zero presence in a particular group. For splicing-factor mutant enrichment, the same statistics were applied, but the samples were grouped into four splicing groups (SRSF2, U2AF1_S34F, U2AF1_Q157P, and SF3B1), along with all other tumors and CD34-positive cells as comparators. Nonsense-mediated decay predictions were extracted from SQANTI3 annotations.

Differentially expressed isoforms were detected using rMATS-long v2.0.1, with parameters `--delta-proportion 0.1` and `--adj-pvalue 0.1`, which also requires average reads per group > 10.0 and transcript CPM >= 5% of gene CPM.

#### Data Portal

The data exploration portal was based on the IsoVis software, with substantial modifications made to enable abundance plotting, optimization to allow for larger datasets, exposing additional transcript information, and providing accessory scripts for preparing data.[39]. The resulting code is freely available at https://github.com/chrisamiller/aml-transcriptome, and the version described in this manuscript has been stably deposited at https://doi.org/10.5281/zenodo.20314851.

### Supplementary Results

#### Read Truncation Patterns

In an attempt to characterize truncated non-full-splice-match reads, we identified "hotspots" with the following characteristics: a) a 100bp window that contained at least 10 premature terminations with characteristic softclipping b) truncated reads represented at least 5% of the reads in that window, c) the sites had a standard deviation of dispersion greater than 10bp (to exclude artifactual pileups at a single hard-to-sequence location). d) does not occur within 200bp of a transcription start site, to exclude the ends of genuine full-length transcript reads. 130,596 of these hotspots were identified across the 71 samples, with a median of 1682 hotspots per sample (range 430 to 6330). Hotspots occurred in a median of 1057 genes per sample, . The GC content of these regions was significantly, but not dramatically, higher than randomly selected genic windows (50.7% vs 46.0%, Wilcoxon rank-sum p<2e-16). They were also not dominated by homopolymer tracts, or other obvious sequence motifs that would suggest specific mechanisms of cDNA-truncating events.

#### More novel transcripts enriched in SRSF2-mutant samples are predicted to undergo NMD

Since we observed an overall shift towards more NMD transcripts in novel isoforms, we wondered whether specific mechanisms of alternative splicing would create higher or lower levels of NMD. When examining known and novel transcripts together, *SRSF2*-mutant samples have non-significant increase levels of NMD transcript expression (0.48% vs 0.44%, Fisher's exact test $p$=0.067). However, when considering only novel transcripts, 13.9% of are predicted to undergo NMD in *SRSF2*-mutant samples compared to 8.8% of of other samples ($p=6.5\times10^{-11}$). In addition, the expression of novel NMD transcripts in *SRSF2* samples is higher than in other samples: in *SRSF2*-mutant samples the mean TPM summed across novel NMD transcripts was 85.49 vs 10.84 in others ($p=6.7\times10^{-14}$). Previous reports have suggested that there was no difference in NMD caused by *SRSF2* mutations, and this remains true when looking at all NMD transcripts combined. However, the increased resolution afforded by this new transcriptome and diverse sample set suggests that there is nuance and biology that has previously been obscured.

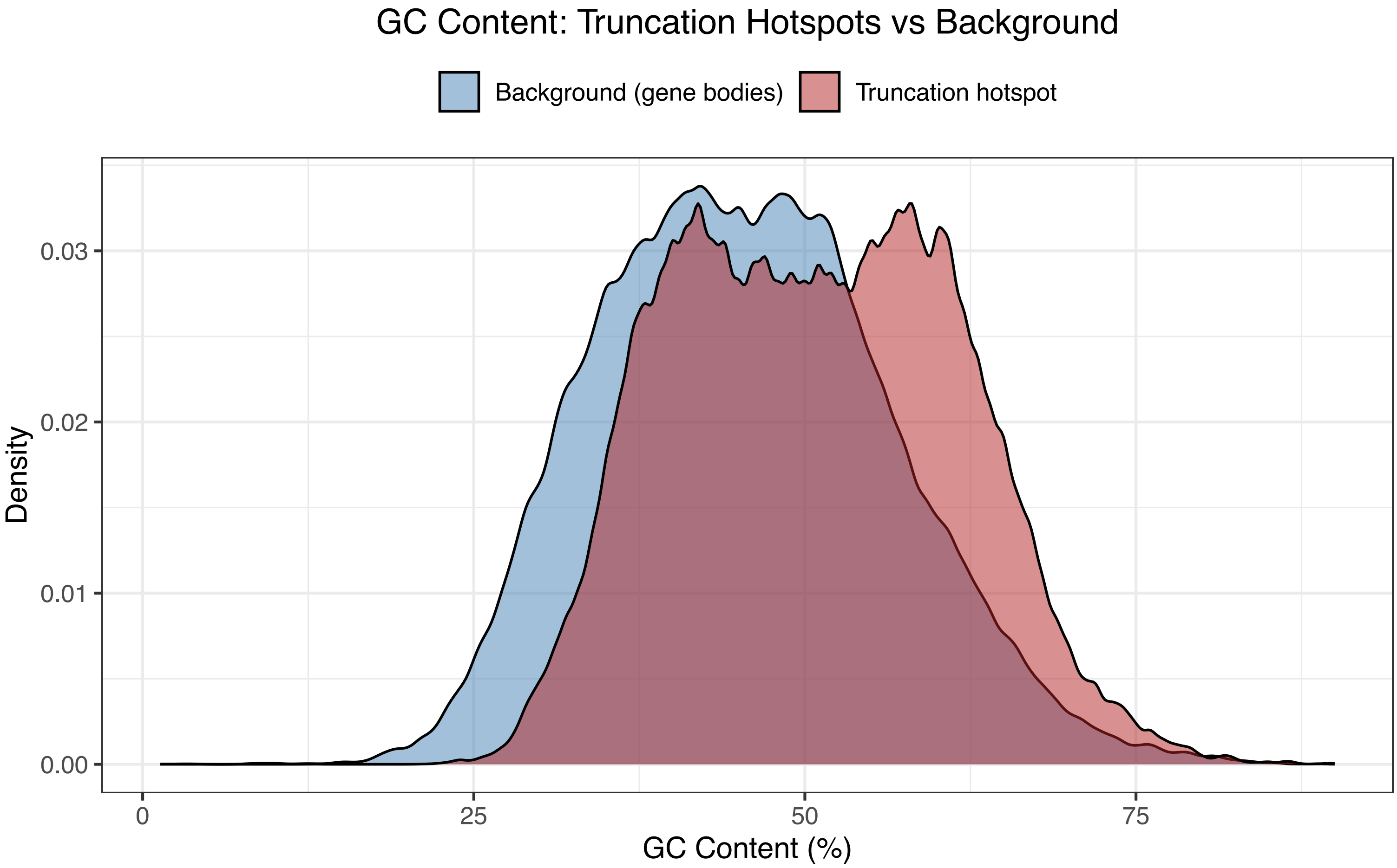


**Figure S1:** Truncation hotspots have significantly higher GC content than randomly sampled sequences from gene bodies with matched length distributions (Wilcoxon rank sum $p < 2.0e-16$)

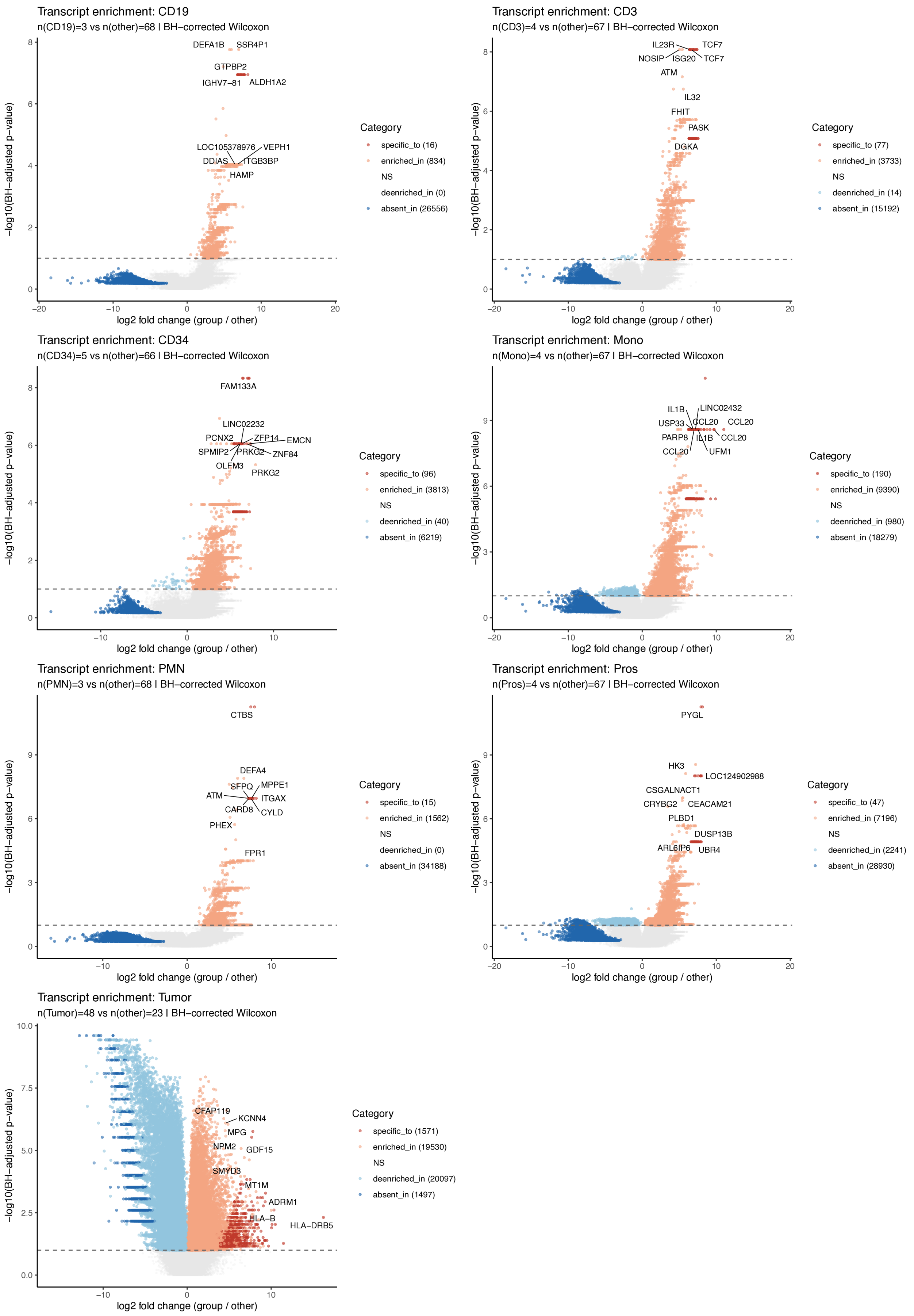


**Figure S2:** Comparisons between specific populations and all other samples, identifying isoforms that were specific to, enriched in, deenriched in, or absent in specific groups

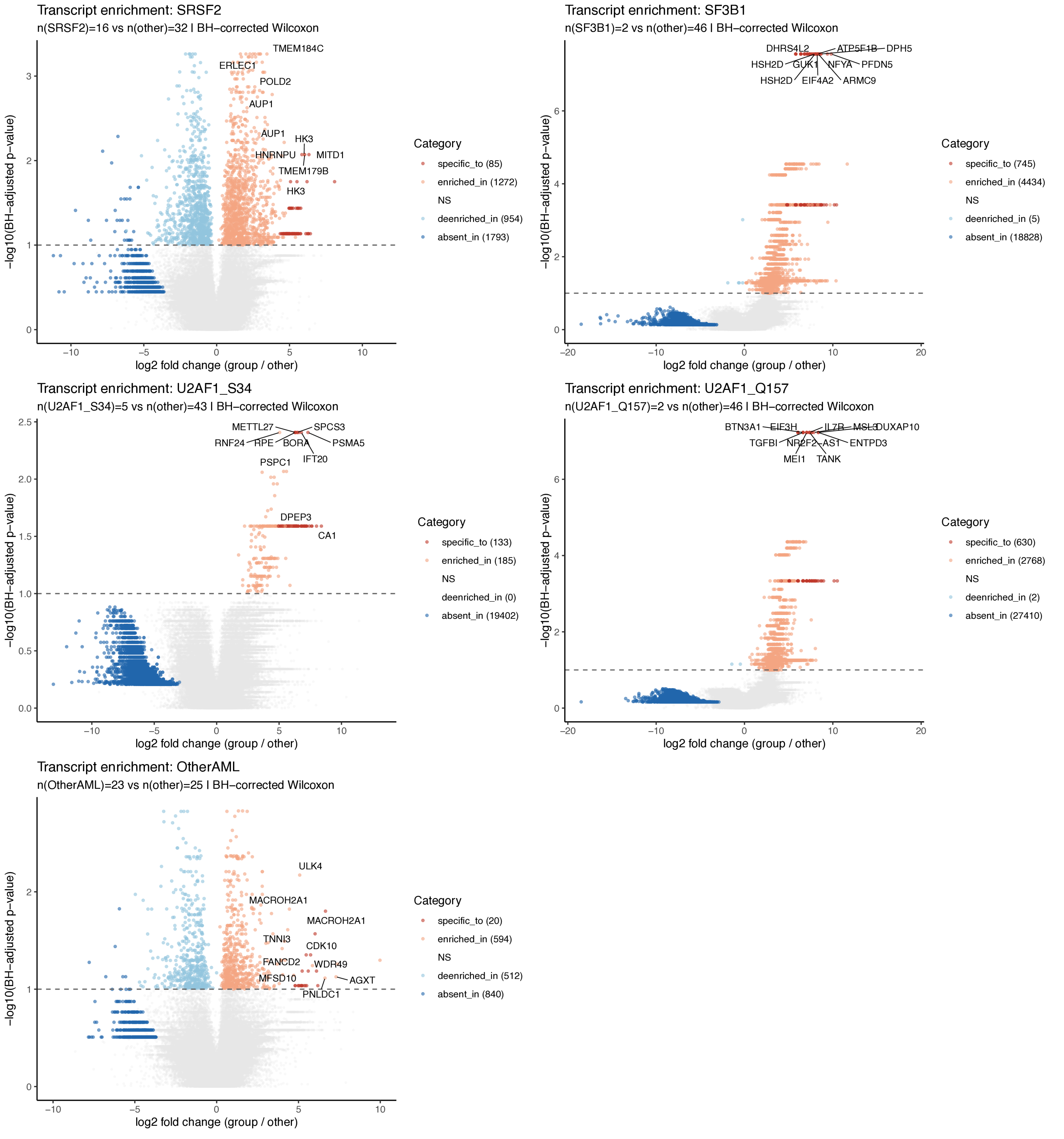


**Figure S3:** Comparisons between specific tumor groups and other tumor samples, identifying isoforms that were specific to, enriched in, deenriched in, or absent in specific groups
